# Generation of iPSC-Derived Human Peripheral Sensory Neurons Releasing Substance P Elicited by TRPV1 Agonists

**DOI:** 10.1101/281675

**Authors:** Marília Zaluar P. Guimarães, Rodrigo De Vecchi, Gabriela Vitória, Jaroslaw K. Sochacki, Bruna S. Paulsen, Igor Lima, Felipe Rodrigues da Silva, Rodrigo F. Madeiro da Costa, Lionel Breton, Stevens K. Rehen

## Abstract

Neural crest stem cells (NCPCs) have been shown to differentiate into various cell types and tissues during embryonic development, including sensory neurons. The few studies addressing the generation of NCPCs and peripheral sensory neurons (PSNs) from human induced Pluripotent Stem Cells (hiPSCs), generated sensory cells without displaying robust activity. Here, we describe an efficient strategy for hiPSCs differentiation into NCPCs and functional PSNs using chemically defined media and factors to achieve efficient differentiation, confirmed by the expression of specific markers. After 10 days hiPSCs differentiated into NCPCs, cells were then maintained in neural induction medium containing defined growth factors for PSNs differentiation, followed by 10 days in neonatal human epidermal keratinocytes-(HEKn-) conditioned medium. We observed a further increase in PSN markers expression and neurites length after conditioned medium treatment. The resulting neurons released substance P (SP) in response to nociceptive agents such as anandamide and resiniferatoxin. Anandamide induced substance P release via activation of TRPV1 and not CB1. Transcriptomic analysis of the PSNs revealed the main dorsal root ganglia (DRG) neuronal markers and a transcriptional profile compatible with C-LTMR. TRPV1 was detected by immunofluorescence and RNA-Seq in multiple experiments. In conclusion, the developed strategy generated PSNs useful for drug screening that could be applied to patient-derived hiPSCs, consisting in a powerful tool to model human diseases *in vitro*.

## 1. Introduction

Human induced pluripotent stem cells (hiPSCs) are used to generate different neuronal types to study biology, pharmacology and to screen for potential new drugs. Developing human peripheral sensory neurons are of great interest, not only to advance pain research aiming at finding new analgesics, but also to the cosmetic industry to develop alternative *in vitro* test methods (Leist et al. 2012, McNeish et al. 2015, Yekkirala et al. 2017). In the field of skin irritation, the activity of Transient Receptor Potential (TRP) channels are of valuable importance, since these channels detect a range of topical nociceptive irritants, such as capsaicin and mustard oil (Nilius and Szallasi, 2014). Capsaicin has been used as a reference in clinical dermatology for skin neurosensitivity and reactivity, with minimal inter individual variation within different ethnic backgrounds (Jourdain et al. 2009, Gueniche et al. 2014).

TRP channels comprise a diverse family of ligand-gated, mostly non-selective, cation channels that are robustly expressed in sensory systems throughout species (Nilius and Szallasi, 2014). Of these, TRPV1 is the most well studied and is considered to be the prototypical TRP channel present in somatosensory neurons (Basbaum et al., 2009). TRPV1 can be directly gated by external molecules such as capsaicin and resiniferatoxin, and also modulated positively or negatively via activation of other receptors and second messenger systems, such as PIP2 hydrolysis and PKC phosphorylation (Julius, 2013). One of the receptors that seem to inhibit TRPV1 activation is the cannabinoid 1 receptor (CB1), also present in somatosensory neurons (Julius and Basbaum, 2001). However, an endogenous agonist of CB1, anandamide, is also a TRPV1 agonist, albeit with an EC50 one order of magnitude higher in the latter (Zygmunt et al., 1999).

Some attempts were made to generate somatosensory neurons from hiPSCs, but so far, these cells have been proved to be difficult to generate robustly. For instance, Chambers et al. (2012) developed a differentiation protocol based on a small molecule screen and successfully obtained neurons that express the main expected markers. Nevertheless, although most neurons exhibited those markers, few showed important functional readouts, such as TRPV1 activation, as only about 1-2% of the cells responded to capsaicin. Other studies were able to demonstrate the expression of canonical peripheral markers, such as BRN3A, peripherin and *β*-Tubulin III, in sensory neurons derived from hiPSCs, but evidence of specific functional activity was still missing (Menendez et al., 2013, Young et al., 2014, Blanchard et al., 2015, Eberhardt et al., 2015). Young et al. (2014) reported measuring TRPV1 activity induced by capsaicin, but only after 6 weeks in medium containing growth factors. Another group showed general electrophysiological activity (*i.e*. action potentials, sodium currents) of human sensory neurons in culture, but did not show responses to capsaicin (Wainger et al., 2015). In the same paper, neurons obtained from mice with a similar method showed robust TRPV1 activity.

Keratinocytes and sensory neurons have an extensive interplay during development and within mature skin. For instance, keratinocytes release neurotrophic factors that induce branching of free nerve endings and neurite outgrowth toward the skin surface (Albers and Davis, 2007). They also release inflammatory mediators involved in responses to tissue damage and hypersensitivity reactions, as well as responses to cold and heat, through receptors of the TRP family of cation channels (Chung et al., 2004). On the other hand, sensory endings do not only transduce sensory signals, but play an active role in the cutaneous metabolism and homeostasis, through the secretion of pro-inflammatory neuropeptides and inflammatory mediators that control vascularization and tissue renewal (Roosterman et al., 2006). Particularly, TRPV1-positive nociceptors may also regulate skin longevity and metabolism, as well as the immune response over aging, as shown in TRPV1 knock-out mice (Riera et al., 2014).

In the glabrous skin, the sensory information is processed by four types of mechanoceptors: Merkel cells, Pacinian corpuscles, Meissner and Ruffini terminations. In the hairy skin, tactile transmission is associated with the lanceolate Aβ-LTMR terminations. Hair follicles are innervated by the longitudinal lanceolate terminations C-LTMR, Aδ-LTMR and Aβ-LTMR. The intercommunication between nerve endings and epidermal keratinocytes occurs through neuropeptides involved in nociceptive and pruriceptive sensitization (Abraira and Ginty, 2013).

Substance P (SP) is a neuropeptide, member of the tachykinin family, synthesized by sensory neurons that emit their extensions from the dorsal root ganglion (DRG) to the more superficial layers of the skin, mediating the communication between peripheral neurons and epidermal keratinocytes (Ribeiro-da-Silva and Hokfelt, 2000). Most of the neurons that release substance P are sensitive to capsaicin, highlighting the importance of TRPV1 expression and sensory neurons-keratinocytes interplay.

Literature analysis reveals the efforts made by several groups to develop more efficient and cost-effective protocols for obtaining functional peripheral neurons from human induced pluripotency stem cells (Menendez et al., 2013, Young et al., 2014 Blanchard et al., 2015, Eberhardt et al., 2015). However, none of these studies have shown results with human cells or investigated the role of the interaction between human epidermal keratinocytes and the transcriptional profile of mature neurons in the presence of factors released by these cells.

In the present work, we developed a new protocol for the direct differentiation of hiPSCs to peripheral sensory neurons in the presence of conditioned medium (CM) obtained from HEKn. These neurons showed some expected functional responses, such as neuropeptide release. The analysis of differential gene expression confirms the transcriptional profile of putative human somatosensory peripheral neurons.

## 2. Material and Methods

### 2.1. Cell Culture

Human induced pluripotent stem cells (hiPSC) (GM23279A from Coriell Institute for Medical Research) were cultured on matrigel-coated plates with mTeSR (Stemcell Technologies, Vancouver, Canada). The cells were checked for pluripotency markers expression such as Nanog, Sox2, Tra-1-60 and Tra-1-81 (data not shown), and were negative for the neuronal markers Islet1, BRN3A, peripherin and TRPV1 (Supplementary Figure 1).

When approximately 60-70% confluence was reached, cells were split using a 0.5 mM EDTA solution (Life Technologies, Rockville, MD, USA). When the same confluence was achieved again, the differentiation was initiated by switching to 3N medium (DMEM-F12, Neurobasal Medium, Glutamax, N2 Supplement, B27 Supplement, non-essential amino acids and β-mercaptoethanol) containing 500 nM LDN-193189 (Stemgen, Cambridge, MA, USA), 10 μM SB431542 (Sigma, St. Louis, MO, USA) and 3 μM CHIR-99021 (Tocris Bioscience, Minneapolis, MN, USA) (d0-d5, Figure 1). Treatment with these Smad pathway inhibitors leads to differentiation of hiPSCs cells toward Neural Crest Progenitor Cells (NCPCs) (Mica et al. 2013) (Figure 1). On day 11 (d11) of differentiation, NCPCs were plated onto Polyornithine (PLO)/Laminin-coated plates and maintained in 3N medium for neuronal differentiation with addition of 10 ng/mL FGF-2 (Life Technologies, PHG0263, Rockville, MD, USA) and 10 ng/mL EGF (Life Technologies, PHG0313, Rockville, MD, USA) for the first 2 days. From D13 to d33 of differentiation, the cells were cultured in 3N medium supplemented with 10 ng/ml BDNF (R&D systems, 248-BD-025,USA), 200 μM ascorbic acid (AA) (Sigma Aldrich, A4034, St. Louis, MO, USA), 10 ng/ml GDNF(R&D Systems, 212-GD-010, USA), 10 ng/ml NGF (R&D Systems, 256-GF-100, USA), 10 ng/ml NT-3 (R&D Systems, 267-N3-025, USA) and 0.5 M cAMP (Sigma-Aldrich, D0260-100MG, St. Louis, MO, USA), resulting in the formation of immature peripheral sensory neurons. After these 35 days, cells could be replated after enzymatic passage with Accutase (Thermo Fisher Scientific, Waltham, MA, USA). Subsequently, neurons were kept for additional 2, 5 or 10 days in conditioned medium from neonatal human epidermal keratinocytes (HEKn), obtained as described in the following session.

**Figure 1:**
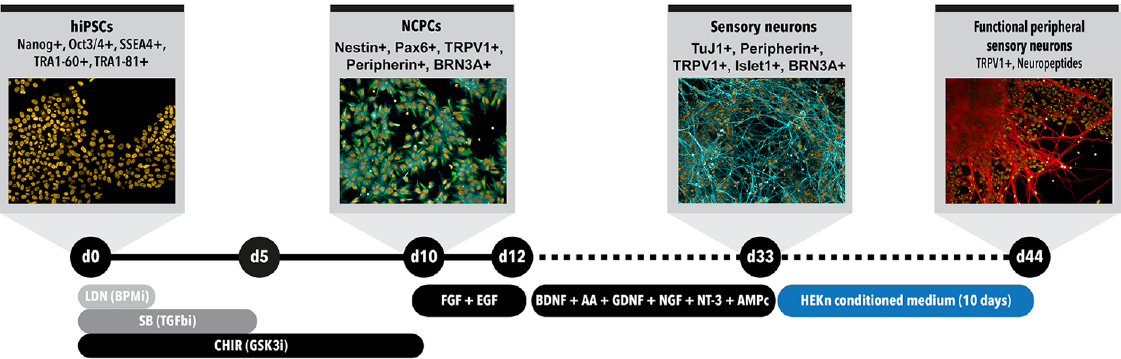
Peripheral sensory neurons differentiation protocol time course. Schematic model of the differentiation protocol to generate sensory neurons from hiPSCs. Day 0 (d0): differentiation starts with hiPSCs presenting all main pluripotency markers; d5: formation of neural tube-like structures; d10: cell commitment to NCPC phenotype; d13: replacement to 2N media supplemented with brain derived neurotrophic factor (BDNF), ascorbic acid (AA), glial derived neurotrophic factor (GDNF), neurotrophin 3 (NT-3) and cyclic AMP (cAMP) to induce NCPC differentiation into sensory neurons; d13-d33: maturation of sensory neurons. d34-d44: treatment with conditioned medium from human neonatal epidermal keratinocytes (HEKn).

### 2.2. HEKn-conditioned medium

Plates were previously treated with gelatin (Sigma, St. Louis, MO, USA) and HEKn (Thermo Fisher Scientific, Waltham, MA, USA) were cultured in Epilife medium (Life Technologies, Rockville, MD, USA). Cells were split when 70% confluence was reached. After 48h conditioning, medium was collected, centrifuged to remove debris and dead cells and was added fresh to the neurons at 75% with 25% of neuronal media.

### 2.3. Immunocytochemistry

NCPCs and sensory neurons were cultured in 96-well plates and fixed with 4% paraformaldehyde, permeabilized with Triton X-100 (0.3%) and blocked with 3% fatty acid-free bovine serum albumin (BSA) (Sigma, St. Louis, MO, USA). Cells were incubated for 2 hours with primary antibodies diluted in 3% BSA. After washing with PBS, fluorescently labeled secondary antibodies were added for 40 minutes in the dark, washed thoroughly with PBS followed by a 5-minute incubation with DAPI (4’,6-diamidino-2-phenylindole) for nuclear staining. After rinsing with PBS and water, 50 μL of glycerol was added as mounting media in each well and the plates were sealed with aluminum sticker before analysis. The primary antibodies used were directed against: nestin (1:100, Sigma, St. Louis, MO, USA), β-tubulin Class III (1:200, Merck-Millipore, Merck Millipore, Darmstadt, Germany), Islet1 (1:1000, Abcam, Cambridge, United Kingdom), TRPV1 (1:1000, Abcam, Cambridge, United Kingdom), BRN3A (1:250, Abcam, Cambridge, United Kingdom) and peripherin (1:250, Santa Cruz Biotechnology, Dallas, TX, USA). Secondary antibodies conjugated with Alexa Fluor 488 and Alexa Fluor 594 (1:400, Life Technologies, Rockville, MD, USA). Images were acquired with a High-Content Screening microscope, Operetta (PerkinElmer, Waltham, MA, USA) using a high numerical aperture 20x objective lens and specific filters for fluorescence excitation and emission. Image analyses were performed using high-content image analysis software Harmony 5.1 (PerkinElmer, Waltham, MA, USA) for morphology analysis and fluorescence quantification. Briefly, for immunostaining quantification and cell counting, cells were segmented by proprietary algorithm (PerkinElmer, Waltham, MA, USA) in a constitutive marker channel (eg. TuJ1 for neurons or Nestin for NCPCs). The segmented areas in each image constituted a template used to measure fluorescence in the same image but in the channels related to the staining of interest. Three independent experiments were performed for immunofluorescence imaging and quantifications. Each marker was probed in triplicate in each experiment and twenty different unbiased microscopic fields were imaged in each replicate. Neurite length measurements were performed using proprietary algorithms as described above, however, TuJ1 staining was used as the primary marker for identifying, segmenting and measuring neuronal projections. The number of cell in each image was counted with DAPI staining and the ratio of TuJ1+ cells and DAPI stained nuclei was calculated. Negative controls for immunocytochemistry were obtained by withholding the primary antibodies and resulted in no staining (Supplementary figures 2 and 3).

### 2.4. ELISA

Substance P (SP) levels were determined using a commercially available ELISA kit (Cayman, Pickerington, OH, USA), following the manufacturer’s instructions. Sensory neurons were plated in 24-well plates at high confluence. After 24 h, cells were rinsed with Hanks’ Balanced Salt Solution (HBSS) buffer, composed of (in mM): 145 NaCl, 5 KCl, 1.2 NaHPO_4_, 1.5 CaCl_2_, 1 MgCl_2_, 10 D-glucose, 5 HEPES, pH 7.4. Following washing, the cells were incubated with different concentrations of TRPV1 agonists for other agents or 45 minutes in the same buffer. When used, antagonists were applied 15 minutes before the agonist addition. Subsequently, the supernatants were collected and immediately assayed with the ELISA kit, following the manufacturer’s instructions. All experiments were performed in triplicate and absorbance levels were measured in a microplate reader.

Anandamide (AEA), rimonabant (SR141716A) and resiniferatoxin were obtained from Cayman Chemical Co. (Michigan, USA) and hydrogen peroxide from Calbiochem.

### 2.5. Transcriptomic analysis

Total RNA from three independent cultures of hiPSC-derived neurons, matured in the absence (EP) or presence of human epidermal keratinocyte conditioned medium (CM) were extracted using QIAGEN miRNAasy kit (Invitrogen, Carlsbad, CA, USA), following the manufacturer’s instructions. RNA concentrations were measured by spectrophotometry using Nanodrop (Thermo Fisher Scientific, Waltham, MA, USA). The integrity of the RNA was analyzed using Bioanalyzer (Agilent Technologies, Santa Clara, CA, USA) and the RNA integrity number (RIN) was obtained by identifying the ribosomal subunits in the samples. Only the samples with RIN greater than 8 were sequenced, with an average coverage of more than 50 million reads per library. RNA sequencing was performed at Life Sciences Core Facility (LaCTAD, State University of Campinas, São Paulo, Brazil) using HiSeq 2500 platform (Illumina) in a paired-end mode (Table 1 in supplementary material).

Quality analysis was performed with Fastqc and Trimmomatic 0.36. Reads passed in the quality control were used to calculate 6transcript expression profile with the software Kallisto 0.43.1 (Bray et al. 2016), using bootstrap 30 and kmer-size 31 with ENSEMBL GRC38 *Homo sapiens* release 88 transcriptome. Kallisto output was used as input in the software Sleuth (Pimentel et al. 2017) to perform the differentially expressed genes (DEG) analysis. Only transcripts with an adjusted p-value<0.01, using Benjamini-Hochberg method, were considered significant. The Venn diagrams were generated using the Interactive Venn tool available at http://www.interactivenn.net/ (Heberle et al. 2015). The transcriptional profile of the PSNs were compared with Genotype-Tissue Expression (GTEx) public data using Heat^*^seq tool (Devailly et al. 2016) available at http://www.heatstarseq.roslin.ed.ac.uk. Correlation heatmap was generated from average RNA-Seq data obtained from six biological replicates of PSNs cultures, comparing with gene expression levels of neuronal and skin tissues. Functional protein association network of hiPSC-derived PSN expressed genes was built using STRING version 10.5, available at https://string-db.org/ (Szklarczyk et al. 2015).

### 2.6. Statistical analysis

Graph results are presented as average ± S.E.M., and differences were tested for significance as indicated in the figure legends. All statistical tests were performed with Graph Pad Prism 6. For transcriptomic evaluation, differentially expressed genes analyses were performed using Sleuth (Pimentel et al., 2017), with qval output equivalent to False Discovery Rate (FDR) with adjusted p-value using the Benjamini-Hochberg test.

## 3. RESULTS

### 3.1. Neural Crest Progenitor Cells (NCPCs) differentiation from human iPS cells

hiPSCs were differentiated into NCPCs after Smad inhibitors treatment for 10 days (d10, Figure 1). Cells were then dissociated, cultured in 96-well plates and characterized for the expression of specific markers. NCPCs were positive for Nestin (Figure 2A, D, G, J and M), TRPV1 (Figure 2B), Peripherin (Figure 2E), BRN3A (Figure 2H), Pax6 (Figure 2K) (Zhang et al. 2010) and negative for Islet1 (Figure 2N). The expression of these markers was quantified as shown in Figure 2P.

**Figure 2:**
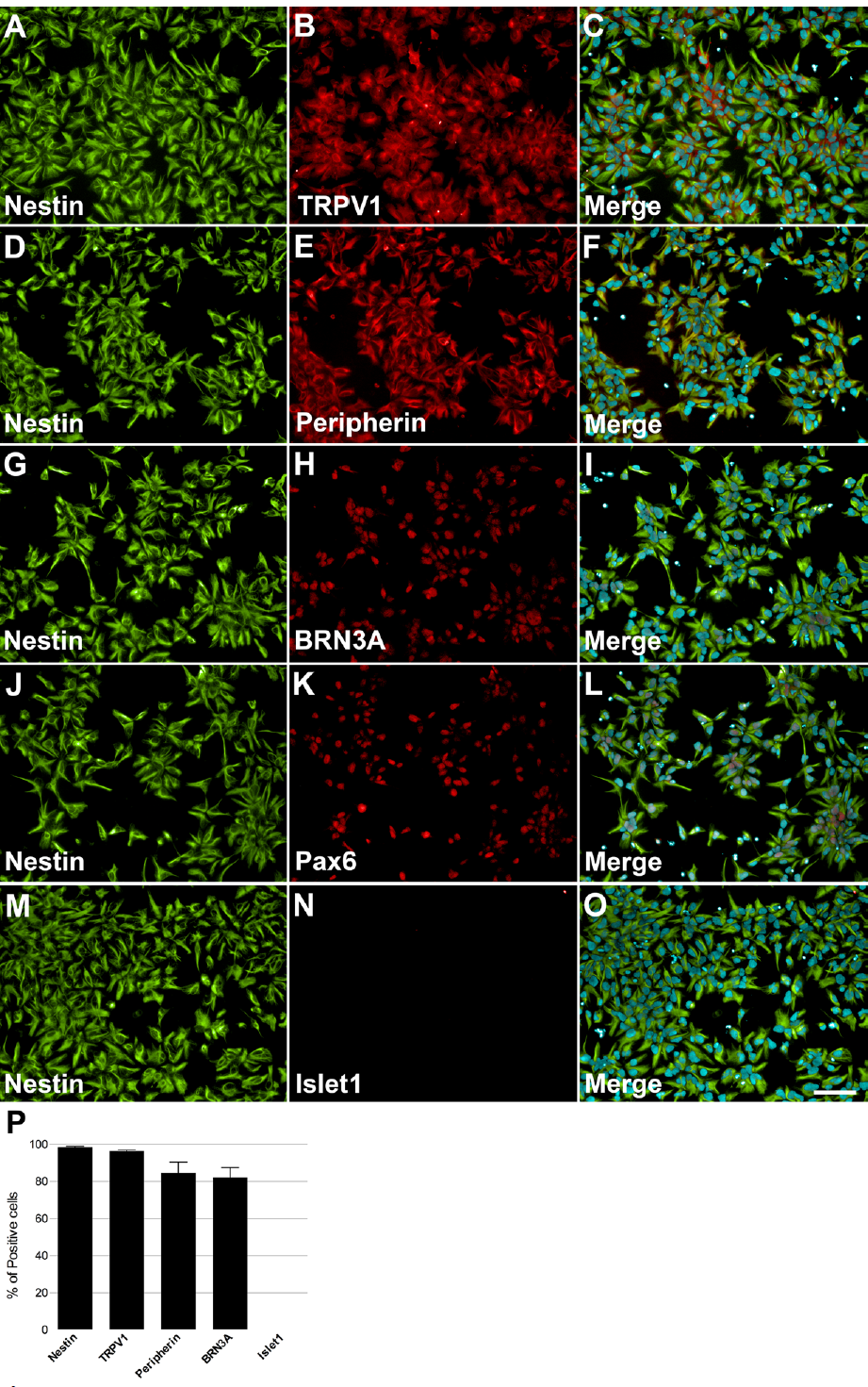
Characterization of NCPCs with specific markers. On day 10, cells presented neural progenitor cell’s morphology and positive staining for Nestin (A, D, G, J and M), TRPV1 (B), Peripherin (E), BRN3A (H), Pax6 (K), and negative for Islet1 (N). Nuclei were stained with DAPI and superimposed with staining for nestin and TRPV1 (C), Peripherin (F), BRN3A (I), Pax6 (L) and Islet1 (O). Calibration bar = 100 μm. Quantification of positive cells for each marker relative to DAPI-stained nuclei, n=2-3 independent experiments (M).

### 3.2. Neural induction

On day 10, NCPCs growth medium was replaced with fibroblast growth factor (FGF-) and epidermal growth factor (EGF-) containing medium for 2 days. Subsequently, cells were plated onto Poly-Ornithine (PLO)/Laminin-coated plates. Cells were maintained in 3N medium with addition of brain derived neurotrophic factor (BDNF), ascorbic acid, glial derived neurotrophic factor (GDNF), nerve growth factor (NGF), neurotrophin-3 (NT-3) and cyclic AMP (cAMP), resulting in the formation of peripheral sensory neurons (Figure 1). Of note, neurons tended to form ganglion-like structures after 7 days in 3N medium, which is also described for mouse and rat primary cultures (Sweetnam et al. 1982). On day 33 (d33) of differentiation, cells were enzymatically detached and replated onto 96-well plates.

Keratinocytes are important in the peripheral sensory neurons maturation process (Mantyh et al. 2011), thus, HEKn conditioned medium (CM) was added to neuronal culture at 75% final concentration for 2, 5 and 10 days before experimental assays. Neuronal maturation was quantified by *β*-Tubulin III staining at these different time points. The neurites emanating from the ganglion-like structures increased robustly over this time, increasing approximately 157% in length after 5 days in culture and 542% after 10 days (Figures 3A, B, C and D). The presence of HEKn CM accounts for a 17.4% increase in cell differentiation, when compared to control cultures maintained in regular 3N medium (Figure 3E).

**Figure 3:**
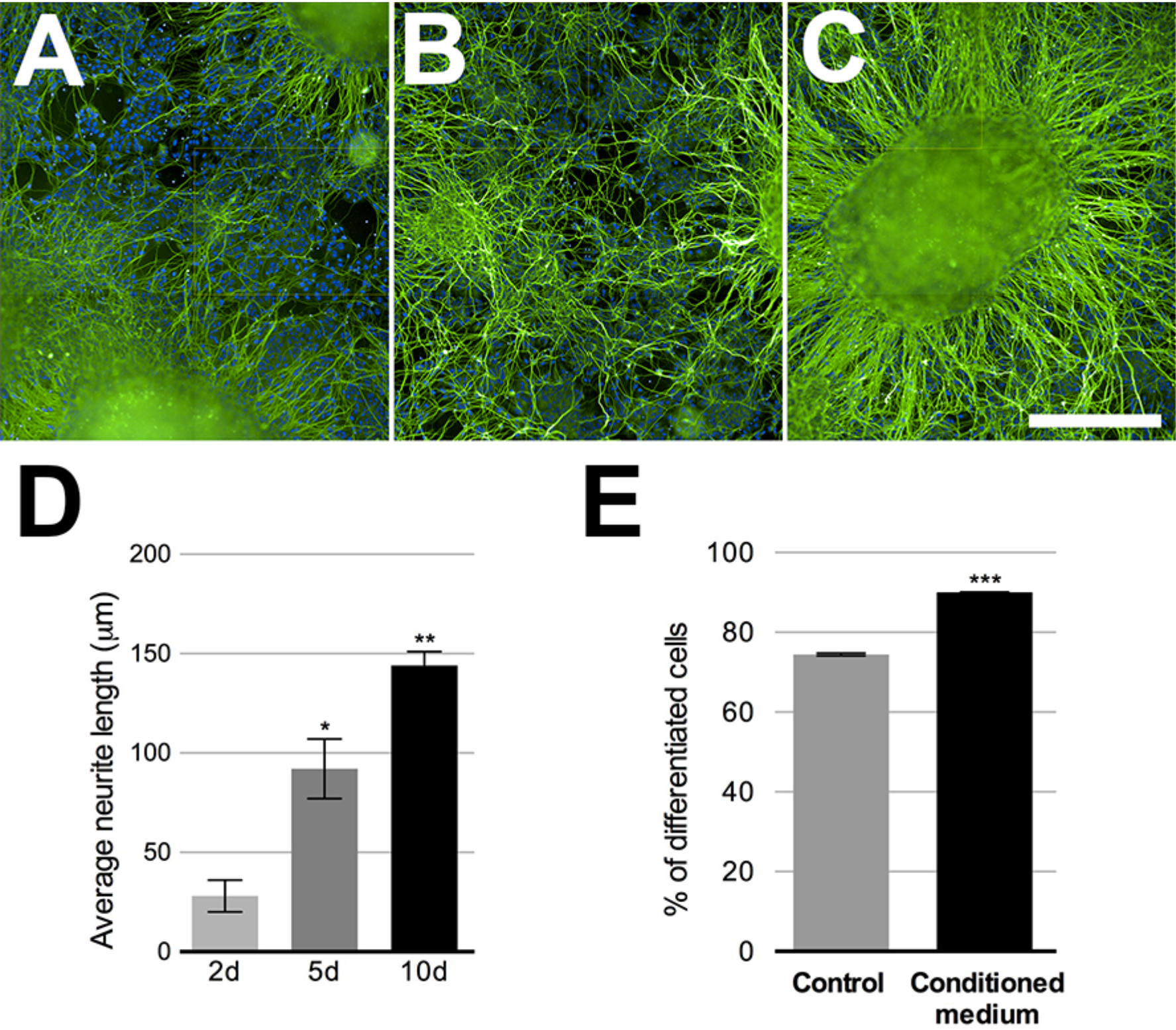
Sensory neurons differentiation. Beta-tubulin class III (green)-stained neurons after 2, 5 or 10 days in neuronal medium conditioned with 75% HEKn medium. Nuclei were stained with DAPI (A-C. Quantification of neurite length in peripheral sensory neurons was based on beta-tubulin Class III (green) immunostaining. Computer assisted image segmentation identified all neurites and measured the total length at each time point. Neurites increased by 328% in length after 5 days and 514% after 10 days in culture with 75% HEKn conditioned medium (D). One-way ANOVA followed by Tukey’s multiple comparison test, ^*^p<0.05 and ^**^ p<0.01 n=3 independent experiments individual experiments from 3 separate differentiations. Percentage of differentiated neurons was increased by HEKn-conditioned medium treatment by 17.4% after 10 days of culture, as measured by βtubulin III-staining (E). Unpaired t test, ^***^p<0.001 n=3 individual experiments from 3 separate differentiations. Scale bar = 100 μm.

There are several markers that make up the distinctive expression profile of somatosensory neurons. We assessed the expression of TRPV1 (Figure 4A-G), Peripherin (Figure 4H-N) and Islet1(Figure 4O-U). All markers showed increased expression over the observed time points (Figures 4G, N and U). TRPV1 increase the most noteworthy, enhancing approximately 5 times. These results suggest that conditioned medium promoted the maturation and growth of sensory neurons, as they exhibited higher expression of differentiation markers, hallmarks of the transition into sensory neurons maturity. The expression of all these markers, except Islet1, was also detected among RNA-Seq transcripts (Figure 6A).

**Figure 4:**
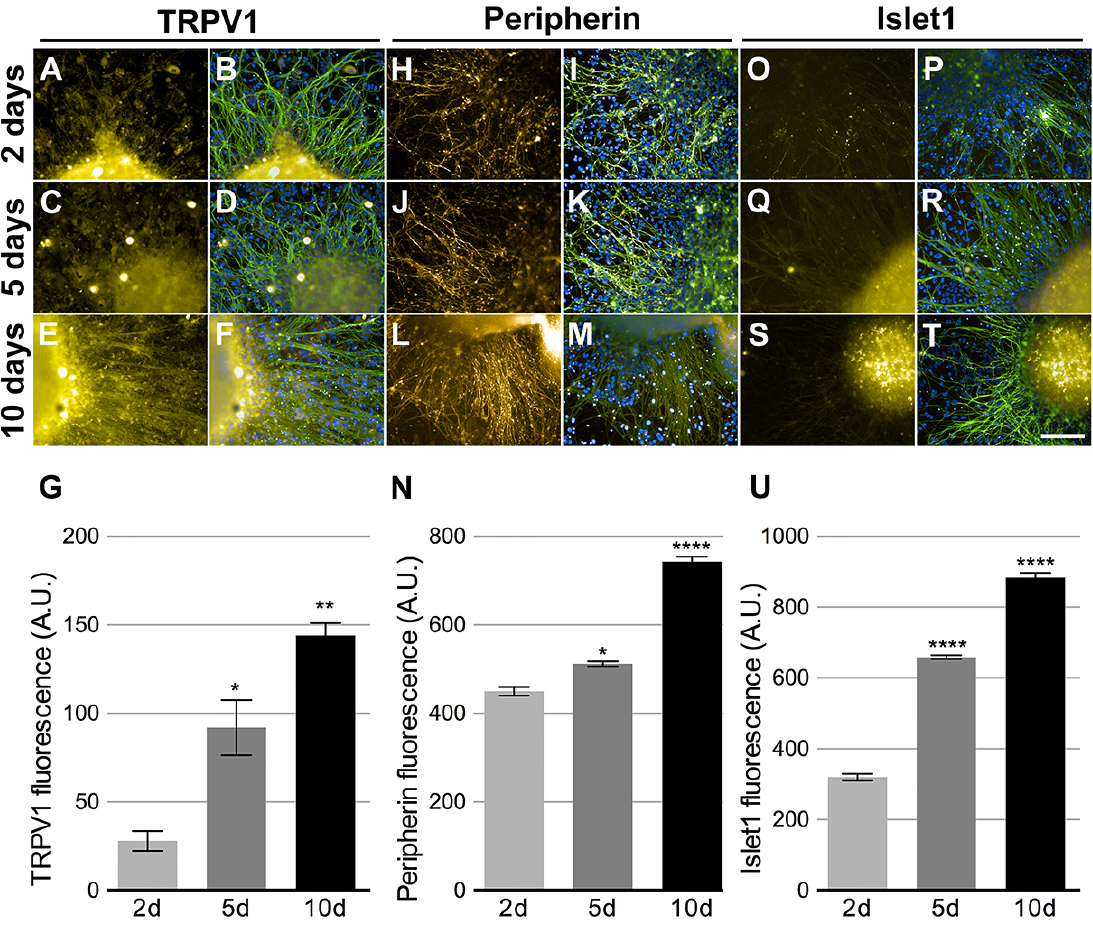
Peripheral sensory neurons maturation. Neurons cultivated in 75% HEKn cells media presented time dependent increase in expression of TRPV1 (A, C, E and G), Peripherin (H, J, L and N) and Islet-1 (O, Q, S and U). Beta-tubulin class III (green) and DAPI (blue) were co-stained in all conditions (B, D, F, L, K, M, P, R and T). TRPV1 fluorescence intensity increased by 328% and 514% after 5 and 10 days, respectively (G); Peripherin showed 118% and 165% increase after 5 and 10 days, respectively (N) and Islet-1 also presented 205% and 276% increase after 5 and 10 days, respectively (U). One-way ANOVA followed by Tukey’s multiple comparison test, ^*^p<0.05, ^**^ p<0.01 and ^****^ p<0.0001, n=3 individual experiments from 3 separate differentiations.

### 3.3. Substance P release caused by nociceptive agents

PSNs respond to irritants, such as capsaicin, by releasing neuropeptides at the epidermal end and glutamate and neuropeptides in the dorsal horn of the spinal cord (Basbaum, Bautista, Scherrer and Julius 2009). To verify whether the sensory neurons were functional with respect to neuropeptide release, after d33 they were plated onto 24-well plates and treated 10 days with HEKn conditioned medium, as described above. Basal Substance P (SP) release was approximately 3 pg/mL/h. This amount was tripled after 300 nM Capsaicin treatment; however, this increase was not statistically significant. A higher concentration of capsaicin was applied (3 μM), but did not lead to an increased release of SP, probably due to TRPV1 desensitization (data not shown). Nevertheless, 300 nM resiniferatoxin effectively increased SP release, as did 20 nM bradykinin (Figure 5A). Interestingly, 100 μM anandamide evoked the highest SP release by the tested TRPV1 agonists with these cells. Moreover, when a high potassium solution (70 mM) was added to depolarize neurons, we obtained an increase in substance P release, though smaller then resiniferatoxin, bradykinin and anandamide (Figure 5A). In addition, the cells responded to 0.25% hydrogen peroxide by releasing the highest amount of SP.

**Figure 5:**
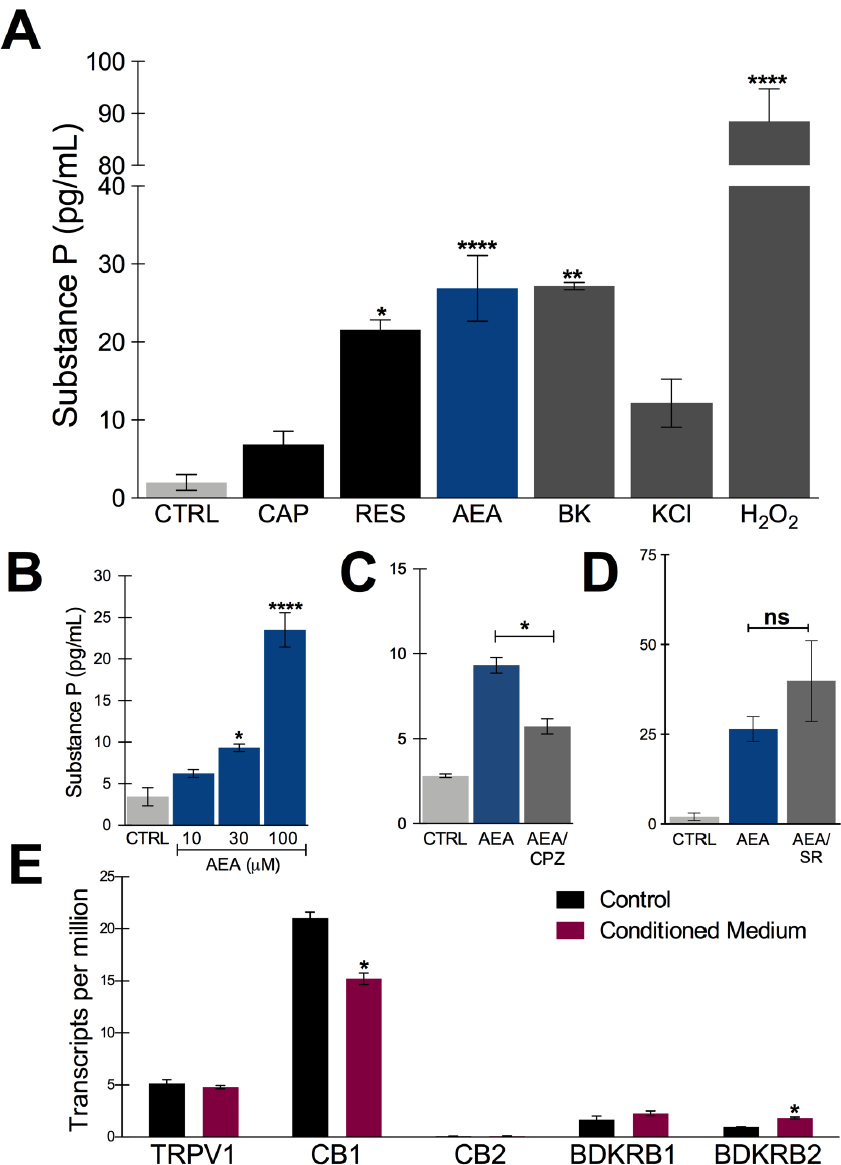
Substance P release after *in vitro* treatment with nociceptive agents. (A) Substance P release measured in the supernatant of sensory neurons culture after incubating for 45 min at 37°C in the presence of ligands. CAP: capsaicin (300 nM); RES: resiniferatoxin (300 nM), AEA: anandamide (100 μM, unless stated otherwise); BK: bradykinin (20 nM); KCl: high potassium solution (70 mM) and H_2_O_2_ (0.25%). (B) concentration-dependent effect of anandamide in substance P release. (C) anandamide effect (30 μM) is partially inhibited by capsazepine (CPZ) (30 μM). (D) anandamide effect (100 μM) is not significantly changed by rimonabant (SR) (10 μM). (E) Transcript levels of the putative receptors involved in substance P release by the tested agonists. TRPV1; transient receptor potential vanilloid 1, CB1; cannabinoid receptor 1, CB2; cannabinoid receptor 2, BDKRD1; bradykinin receptor 1 and BDKRD2; bradykinin receptor 2. One Way ANOVA followed by Tukey’s multiple comparison test, ^*^p<0.05, ^**^ p<0.01 and ^****^ p<0.0001. n=2-3 individual experiments from 3 separate differentiations.

The anandamide effect, albeit at relatively high concentrations, was concentration-dependent in a range compatible with TRPV1 agonism rather than CB1 activation (Figure 5B). Furthermore, AEA-increase of SP release was partially inhibited by capsazepine, a TRPV1-selective antagonist (Figure 5C). Rimonabant, a CB1-selective antagonist did not block the AEA effect, on the contrary, it tended to increase it (Figure 5D). These results indicate that functional nociceptive responses are present in these neurons, particularly TRPV1-mediated ones. The expression of the potential receptors involved in these functional responses are plotted in Figure 5E and include TRPV1, CB1 and CB2, and both bradykinin receptors, BDKRD1 and BDKRD2. Interestingly, CB2 was detected in negligible amounts.

### 3.4. Transcriptomic analyses

RNA-Seq was used to determine the transcriptional profile of the cells obtained using the differentiation protocol aforementioned. RNA from control PSN (PSN-C) and treated with HEKn-conditioned media (PSN-CM) during maturation in culture have both shown expression of all main expected neuronal markers (Figure 6A, B). Transcripts coding the same markers that were detected by immunostaining, such as Nestin (NES), Pax6 (PAX6), Peripherin (PRHP), BRN3A (POU4F1). TRPV1 was the most expressed TRP channel of sensory neurons, confirming the immunofluorescence staining. Also, the presence of the most expressed neuronal marker genes like PIEZO2 (piezo type mechanosensitive ion channel component 2), TH (tyrosine hydroxylase), TUBB3 (tubulin beta 3 class III), Neurotrophic receptor tyrosinase 2 (NTRK2, also known as TrkB), SCN3A and SCN3B (Nav 1.3 or sodium voltage-gated channel alpha and beta subunit 3) suggest that the obtained cell type has a C-Low Threshold mechanoreceptor (C-LTMR) profile, EGF (epidermal growth factor), RUNX1T1 (RUNX1 translocation partner 1), CALCB (CGRP receptor component), LDHB (lactate dehydrogenase B), MDK (midkine, neurite growth-promoting factor 2), MSN (moesin) and P2RX3, P2RX4 (purinergic receptor P2X ligand-gated ion channel 3 and 4), were found at different levels in both PSN-C and PSN-CM transcriptional fingerprints, in three independent cultures per condition (Figure 6A and Table 1).

**Table 1:**
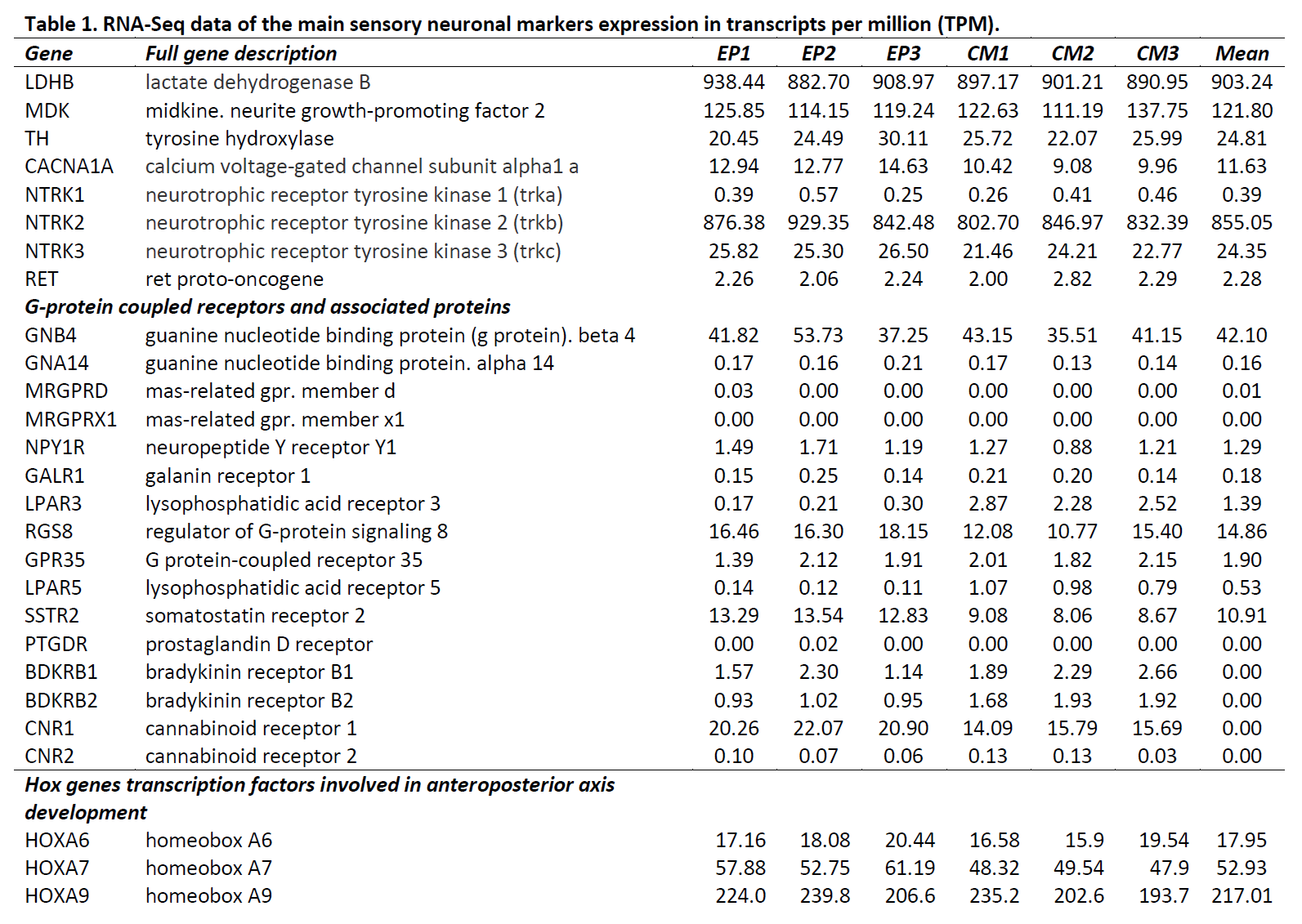
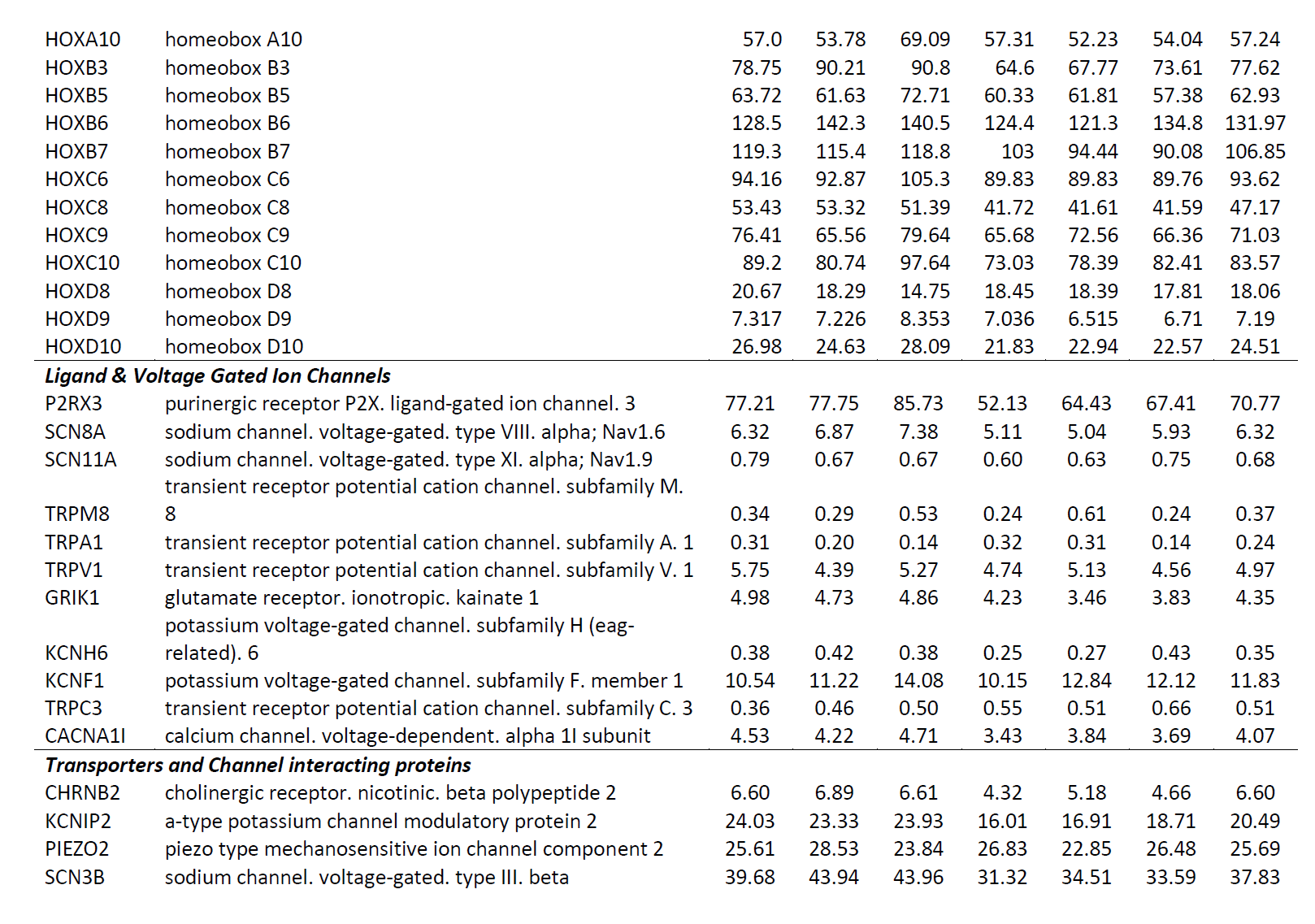
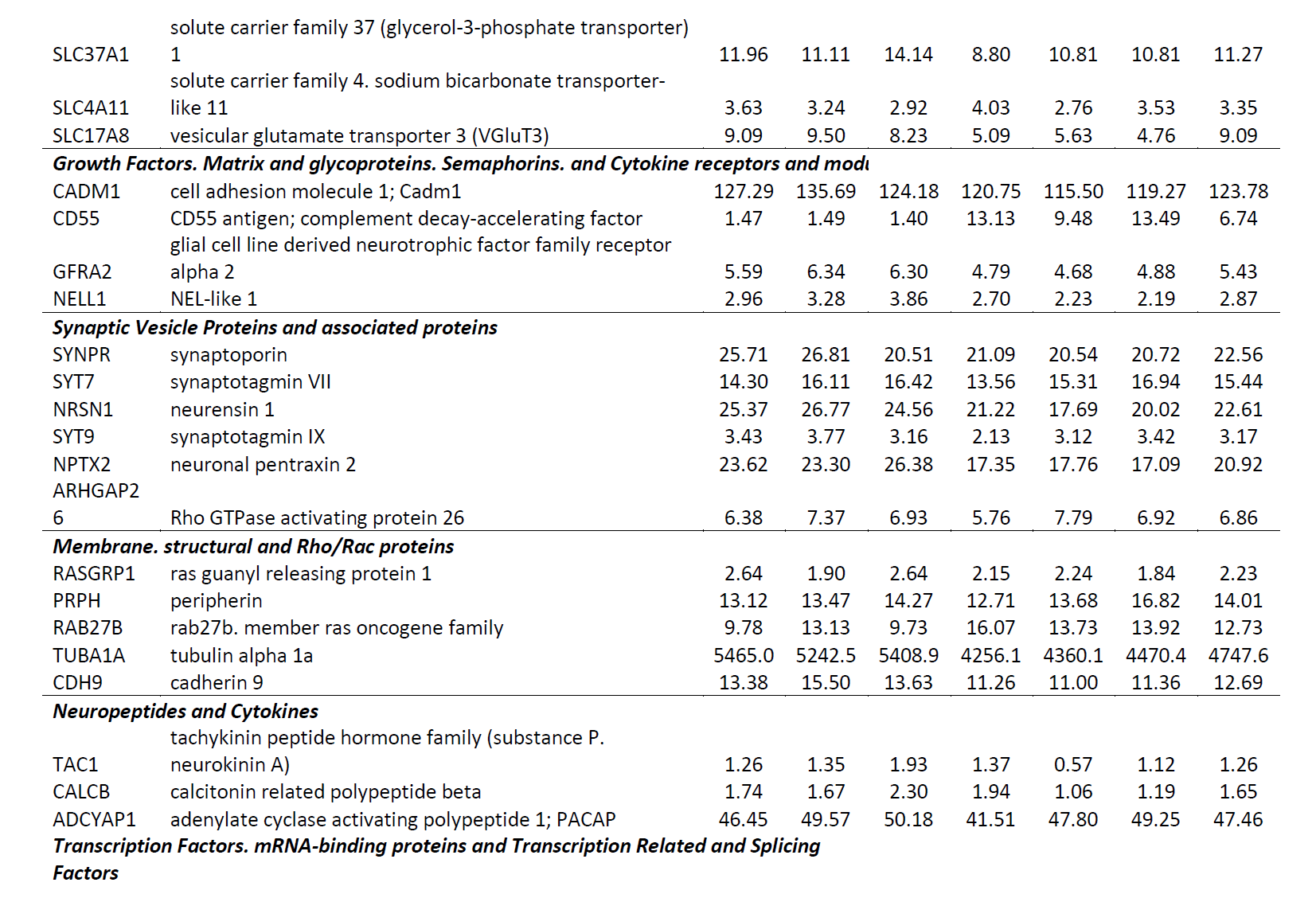
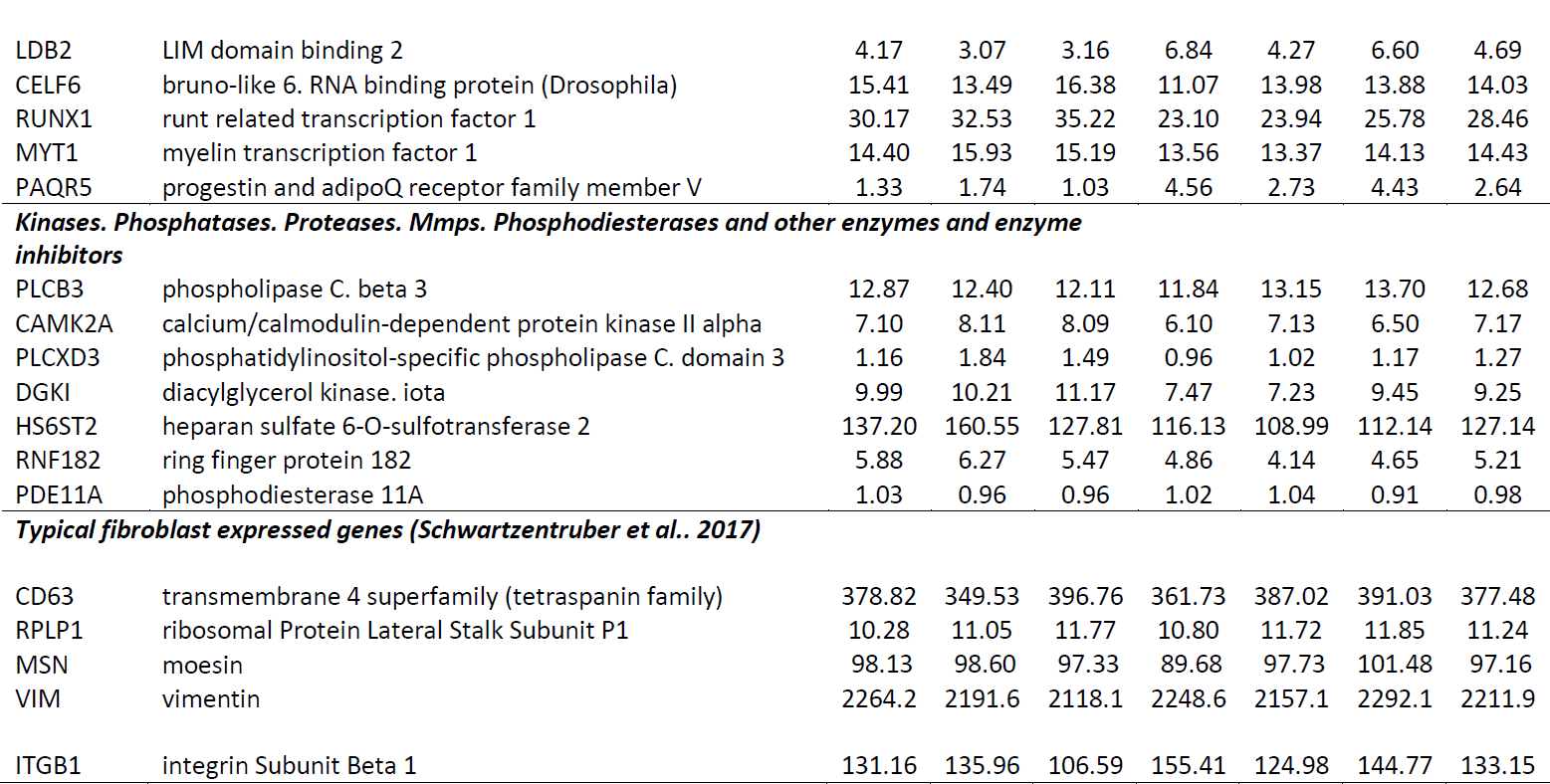
RNA-Seq data of the main sensory neuronal markers expression in transcripts per million (TPM).

**Figure 6:**
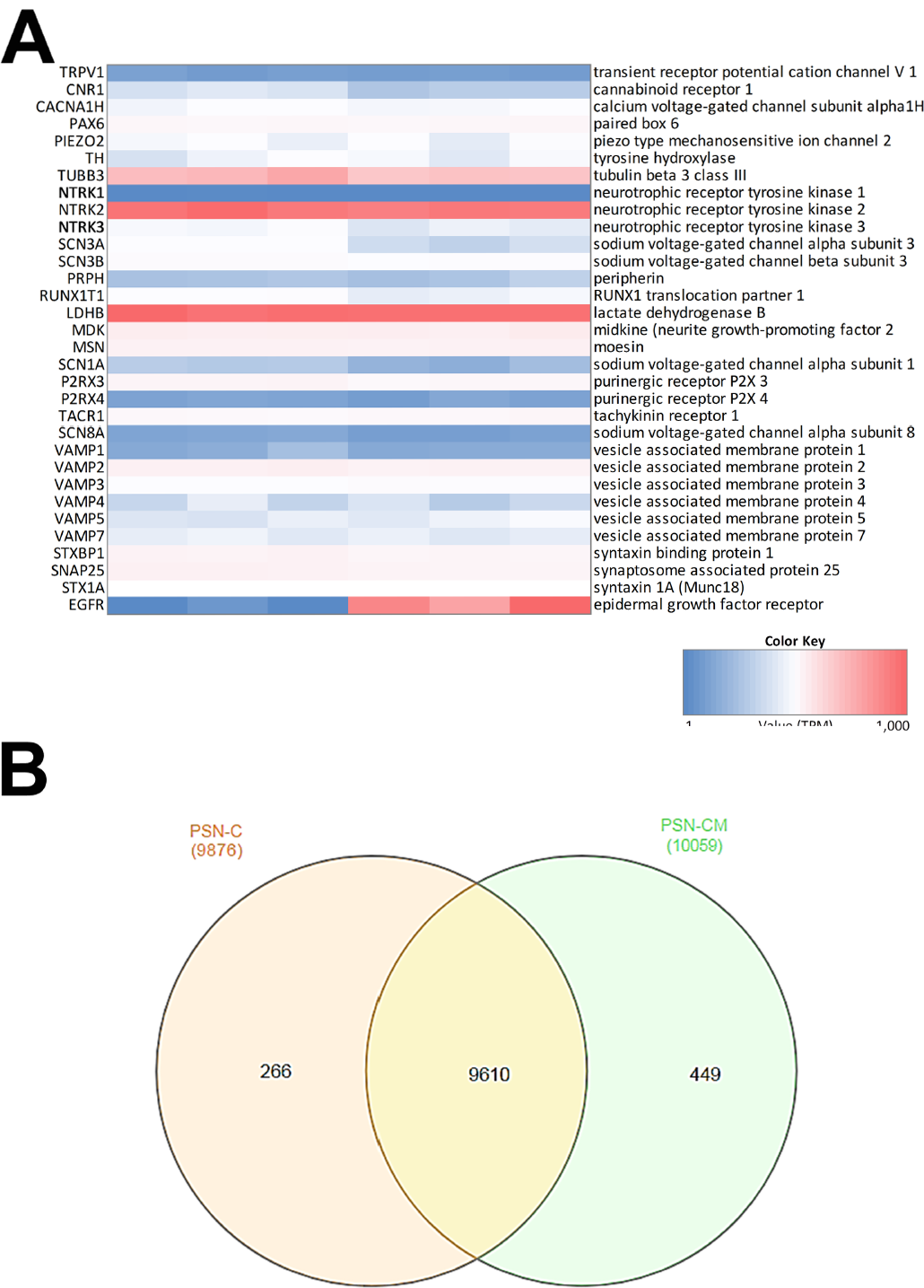
Neuronal markers expressed and regulated in hiPSC-derived peripheral sensory neurons. (A). Heatmap (blue: low expression, red: highly expressed) of transcripts related to neuronal marker genes expressed in cultures of control PSN (PSN-C) and PSN treated with HEK-conditioned media (PSN-CM), obtained from RNA-Seq data (B). Transcripts coding marker genes Nestin (NES), Peripherin (PRHP), BRN3a (POU4F1). Moreover, mRNA encoding the cannabinoid receptor 1 (CNR1) were found in higher levels compared to TRPV1, which was also expressed but in lower levels. PIEZO2 (piezo type mechanosensitive ion channel component 2), TH (tyrosine hydroxylase), TUBB3 (tubulin beta 3 class III), Neurotrophic receptor tyrosinase 2 (NTRK2, also known as TrkB), SCN3A and SCN3B (Nav 1.3 or sodium voltage-gated channel alpha and beta subunit 3), EGF (epidermal growth factor), RUNX1T1 (RUNX1 translocation partner 1), CRCP (CGRP receptor component), LDHB (lactate dehydrogenase B), MDK (midkine, neurite growth-promoting factor 2), MSN (moesin) and P2RX3, P2RX4 (purinergic receptor P2X ligand-gated ion channel 3 and 4) were highly expressed in all cultures. (B) Venn diagrams were generated using InteractiVenn tool showing unique and shared genes expressed in PSN-C and PSN-CM (treated with HEKn-CM).

Venn diagrams (Figure 6B) show about 10,325 genes significantly expressed (TPM>10), with 9,610 genes expressed in both conditions, 266 expressed only in PSN-C and 449 genes exclusively expressed in PSN-CM condition, which could be interesting targets to be investigated to increase PSN maturation. Most of the transcripts that changed were downregulated after the treatment with HEK-CM (**Supplementary Figure 5).** Moreover, mRNA encoding the cannabinoid receptor 1 (CNR1) was found in higher levels compared to TRPV1 (Figure 5E). This could mean that the AEA-induced SP release via TRPV1 could be underestimated because CB1 is also stimulated by anandamide and might inhibit TRPV1. In fact, neurons pretreated with rimonabant, a CB1-selective antagonist, show a tendency to increase SP release with anandamide (Figure 5D). Interleukin 1 α and β (IL1α, IL1β) transcripts were found only in PSN-CM cultures. Principal component analysis (PCA) was performed using ClustVis and provided an accurate verification of the applied method, as the first principle component clearly separated the samples by condition (EP vs CM) and the second principle component separated the samples by independent PSN cultures differentiation (Supplementary Figure 4).

The transcriptome comparison between PSN and GTEx RNA-Seq dataset from human tissues shows higher scaled correlation with brain tissues like hypothalamus, amygdala, frontal cortex, hippocampus (scaled correlation 0.950; 0.936; 0.935, and 0.935 respectively) (Figure 7), suggesting that PSN present a transcriptional profile closer to neuronal than other human tissues or skin cells, like fibroblasts, which were the source of the cells that were reprogramed to generate the hiPSC used in this study.

**Figure 7:**
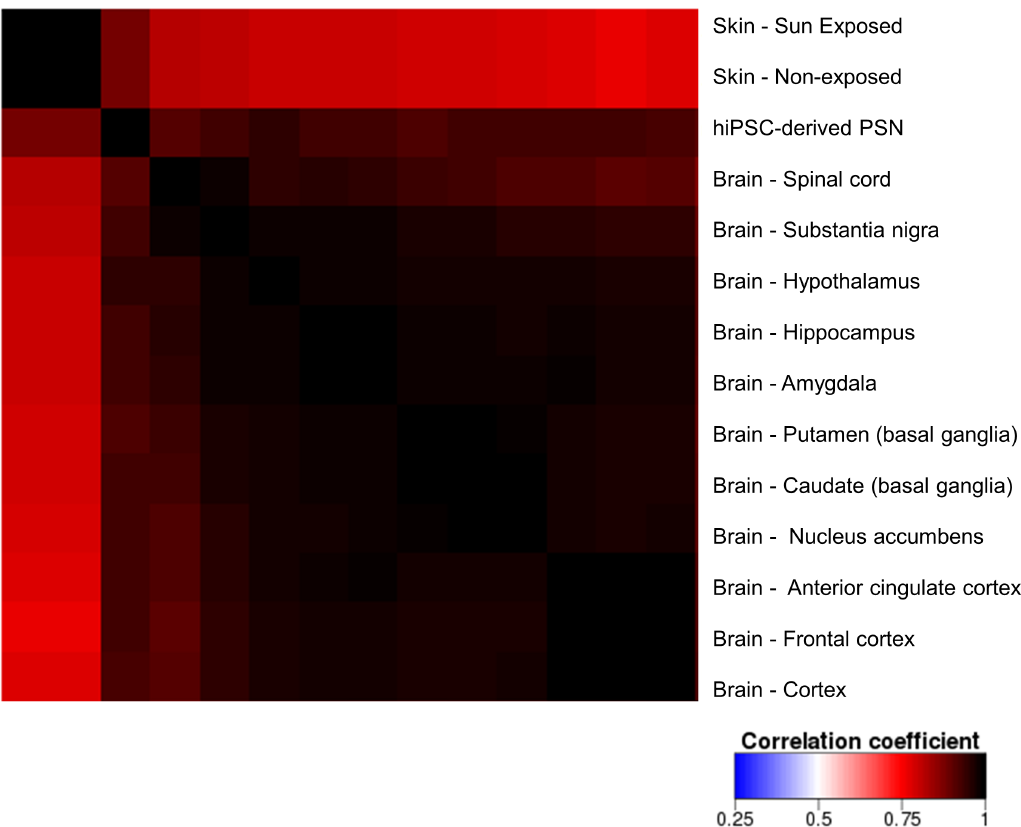
Correlation heatmap comparing PSN transcriptional profile with human GTEx gene expression data of neuronal and skin tissues. Heatmap was generated using Heat^*^seq tool^43^ using average RNA-Seq data from six distinct biological replicates of PSNs cultures, showing higher scaled correlation with neuronal human tissues such as hypothalamus, amygdala, frontal cortex, hippocampus (scaled correlation 0.950; 0.936; 0.935, and 0.935 respectively).

With the RNA-Seq it is possible to identify transcripts uniquely expressed in neuronal subtypes, directly related to functionality and the identification of neuronal types. The relevance of these markers was previously validated *in vivo* with identified population-specific genes in high fraction of positive cells (Usoskin et al. 2015). Neuronal population expressing PIEZO2 and an exclusive C type Low-Threshold MechanoReceptor (C-LTMRs) Slc17a8, codifying the vesicular glutamate transporter 3 (Vglut3), involved in mechanical pain and pleasant touch (Seal et al. 2009, Coste et al. 2010, Li et al. 2011, Abraira and Ginty 2013).

The transcriptomic analysis confirmed the expression of the main neuronal markers expected for somatossensorial DRG neurons, with predominantly transcriptional profile compatible with C-Low Threshold Mechanoreceptors (C-LTMR) showed in Figure 7.

## 4. Discussion

According to literature, several groups have reported generation of peripheral sensory neurons from pluripotent stem cells. However, none was able to show robust TRPV1 activity. Chambers *et al.* (2012) showed TRPV1 expression via quantitative PCR and correlated with only 1-2% of the cells responding to capsaicin. Even the responding cells seem to exhibit a very low activity. Others showed electrophysiological activity of human sensory neurons directly reprogrammed from fibroblasts, but not capsaicin-elicited signals (Wainger et al., 2015). In the same work, mouse neurons obtained with a similar method showed robust TRPV1 activity. One could speculate that the authors were not able to detect TRPV1 activity in human neurons consistently as well. Another work reported detecting TRPV1 activity induced by capsaicin after 6 weeks in media containing growth factors, but did not further describe this response (Young et al., 2014). PSNs generated by other groups that did not demonstrate TRPV1 activity, but focused on sodium currents and action potentials (Menendez et al. 2011, Lee et al. 2012, Reinhardt et al. 2013). To our knowledge, this is the first report of human PSN obtained by any method shown to release SP in response to nocifensive agents.

TRPV1 was detected by immunostaing in ˜ 90% of neurons, but the the RNA levels and substance P release induced by TRPV1 agonists were relatively small. One hypothesis to explain this discrepancy between the TRPV1 staining and function is that, although present, TRPV1 does not reach full activity due to modulation of the presence of this channel on the cell surface, via interaction with other proteins (Figure 8). For instance, CB1 (CNR1) is overexpressed (Figure 5E) relatively to TRPV1. This overexpression could explain the low detectable activity of TRPV1, as CB1 is known to negatively modulate this channel through dephosphorylation (Figure 8) (Ross, 2003, Yang et al., 2013) (Figure 8). Accordingly, acute pretreatment with a selective CB1 inverse agonist tended to augment AEA-induced SP release, presumably through reduced inhibition of TRPV1. However, it remains to be stablished whether chronic suppression of CB1 activity could effectively enhance TRPV1 function.

**Figure 8:**
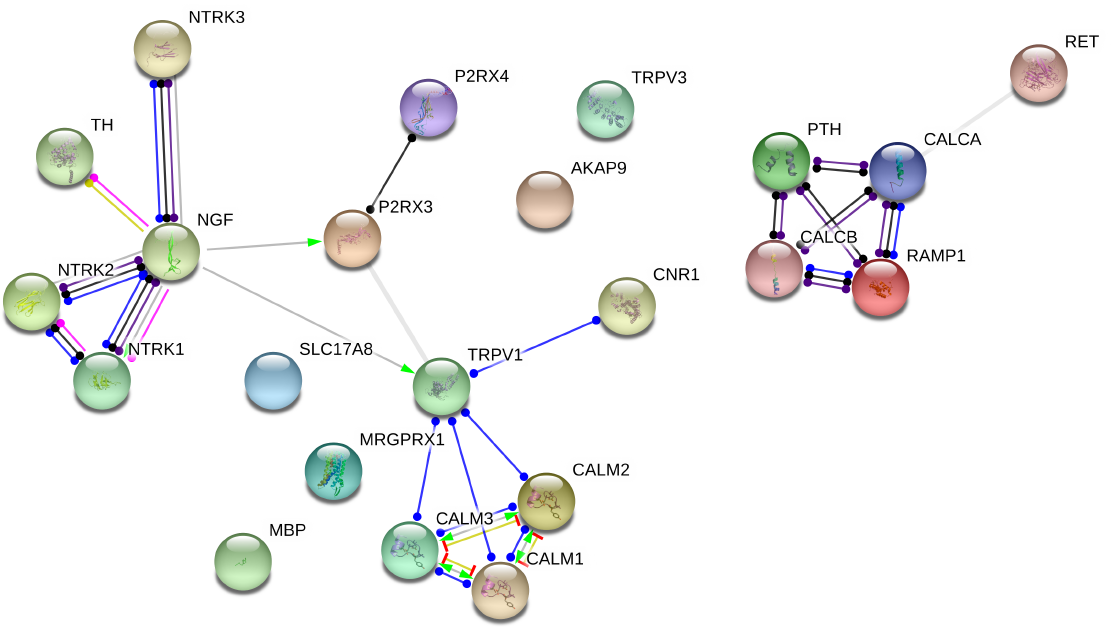
Functional protein association network of hiPSC-derived PSN expressed genes with TRPV1. Hierarchical clustering network of related genes to TRPV1 modulation shows the relationship of TRPV1 with P2RX3, CB1 receptor, CALM 1, 2 and 3.

In addition, in the case of sensory neurons expressing TRPV1 channels, these proteins are packaged into CGRP- or SP- and VAMP1-containing vesicles and delivered to the plasma membrane, involving the formation of SNARE complexes composed of SNAP-25, syntaxin 1, and VAMP1, as well as Munc18–1 (Meng et al. 2016). This delivery can be enhanced by several factors, including TNFα. Since SP release was modestly elicited by capsaicin, it is possible that this aspect of nociceptor development is not completely mature at the differentiation level presented here and therefore one could speculate that TRPV1 is not adequately being shipped to the cell surface.

Furthermore, other proteins known to negatively modulate TRPV1 function that actually are binding partners are calmodulins, which are very highly expressed (data not shown and Figure 8) (Ho et al., 2012). Again, by negatively modulating the levels of these proteins, a more robust TRPV1 activity could arise.

The fact that Young *et al.* (2014) only observed TRPV1 activity after 6 weeks in the presence of growth factors corroborates this notion. However, maintaining cells for another 2 weeks *in vitro* does not guarantee that this activity would be satisfactory in the end, as mentioned. Therefore, we must consider additional mechanisms to enrich the neuronal population in TRPV1+/TrKA+/CGRP+ cells. First, the amount of NGF necessary might be higher, or perhaps a gradient of this neurotrophin might be necessary. NGF is present in high concentrations within the epidermis, where nerve fibers with free endings (i.e. nociceptor fibers) will project (Davies et al., 1987). In murine models, it is known that several steps are necessary to differentiate TrkA+/Met+/CGRP+ neurons, the ones with putative TRPV1 activity (Lallemend and Ernfors, 2012). Our transcriptomic results show a high expression of TrkB, which was reported to be present at high levels in a subset of murine medium-diameter DRG neurons (Li et al., 2011). Although individual cell gene expression variability is influenced by differentiation conditions, our PSN expressed the five sensory-neuronal genes marker genes, except SCN9A (Schwartzentruber et al. 2017). Tyrosine hydroxylase (TH), which catalyzes the production of L-DOPA from tyrosine in the catecholamine biosynthesis pathway, is a defining feature of Low Threshold Mechanoreceptors (LTMRs) in adult DRGs (Brumovsky et al., 2006) and was found to be also highly expressed in hiPSCs-generated PSN. The presence of many individual transcripts characteristic of DRG and somatosensory neurons was confirmed. These include RET, TH, PRPH and LDHB. However, some specific LTMR transcripts were not detected, such as VGLUT3, Nav 1.8 and TRPA1. PIEZO2 and GFRA2, markers of LTMR were expressed and P2RX3, a purinergic ion channel expressed in C-fiber non-peptidergic neurons was also detected (Yin et al., 2016) as shown in Table 1. CACNA1H was highly expressed, being a voltage-gated calcium channel (CaV) subtype 3.2, uniquely expressed in unmyelinated C-LTMR (Reynders et al., 2015). Together with CACNAH1, the high expression of TrkB, TH and LDHB suggest the PSN generated here have a LTMRs profile of peripheral neurons involved in sensory perception (Usoskin et al., 2015). Our RNA-Seq results suggest that the hiPSC-derived PSN present a transcriptional profile compatible with C-LTMR predominantly (Supplementary Figure 6). However, to precisely define the proportion of this neuronal type in the culture further investigations have to be performed using single cell RNA sequencing.

In summary, although some reports have described the generation of human peripheral sensory neurons, none have successfully demonstrated a robust and useful TRPV1 activation. The present work shows detectable TRPV1 activity, the release of SP mediated by resiniferatoxin and anandamide that could be used for screening of nociceptive agents and possibly analgesics. The transcriptional data obtained from RNA-Seq show the main neuronal marker genes expressed, in different levels, in independent PSN cultures. The strategy of treating PSN with CM to investigate whether it could increase cell maturity and TRPV1 functionality had opposed outcomes. Although neurite growth increased, which is an indication of neuronal maturity, there were no conclusive changes in relevant neuronal markers expression in RNA-Seq data, although TRPV1 immunostaining increased. Nevertheless, as TRPV1 activity would be useful for the purpose of screening for new nociceptive/irritant agents, it is appealing to further improve the robustness of this activity and to develop reliable *in vitro* models using these differentiated human neurons.

## Data availability

The datasets generated during and/or analyzed during the current study are available from the corresponding author on reasonable request.

### Acknowledgements

This work was supported by the Brazilian Development Bank (BNDES), Funding Authority for Studies and Projects (FINEP), Research Support Foundation of the State of Rio de Janeiro (FAPERJ), National Council for Scientific and Technological Development (CNPq) and L’Oréal Research and Innovation.

We thank the staff of the Life Sciences Core Facility (LaCTAD) from State University of Campinas (UNICAMP), for the RNA-Seq and support on the Bioinformatics analysis. The authors also wish to thank Dr. Claudia Benjamim and Julio Sharfstein for providing bradykinin and Fernanda Motta and Gabriel Ferraz for the help with calcium experiments. In addition, the authors are thankful for all contribution of Dr. Fabiana Munhoz and for the helpful comments of Aurelia Del Bufalo, before and during the preparation of this manuscript, respectively.

## Author contributions

Conceived and designed the experiments: MZPG RDV JKS BSP RM LB SKR

Performed the experiments: MZPG RDV GV JKS BSP IL RM

Analyzed the data: MZPG RDV BSP RM FRS

Wrote the paper: MZPG RDV BSP RM FRS LB SKR

---

**Supplementary Figure 1:**
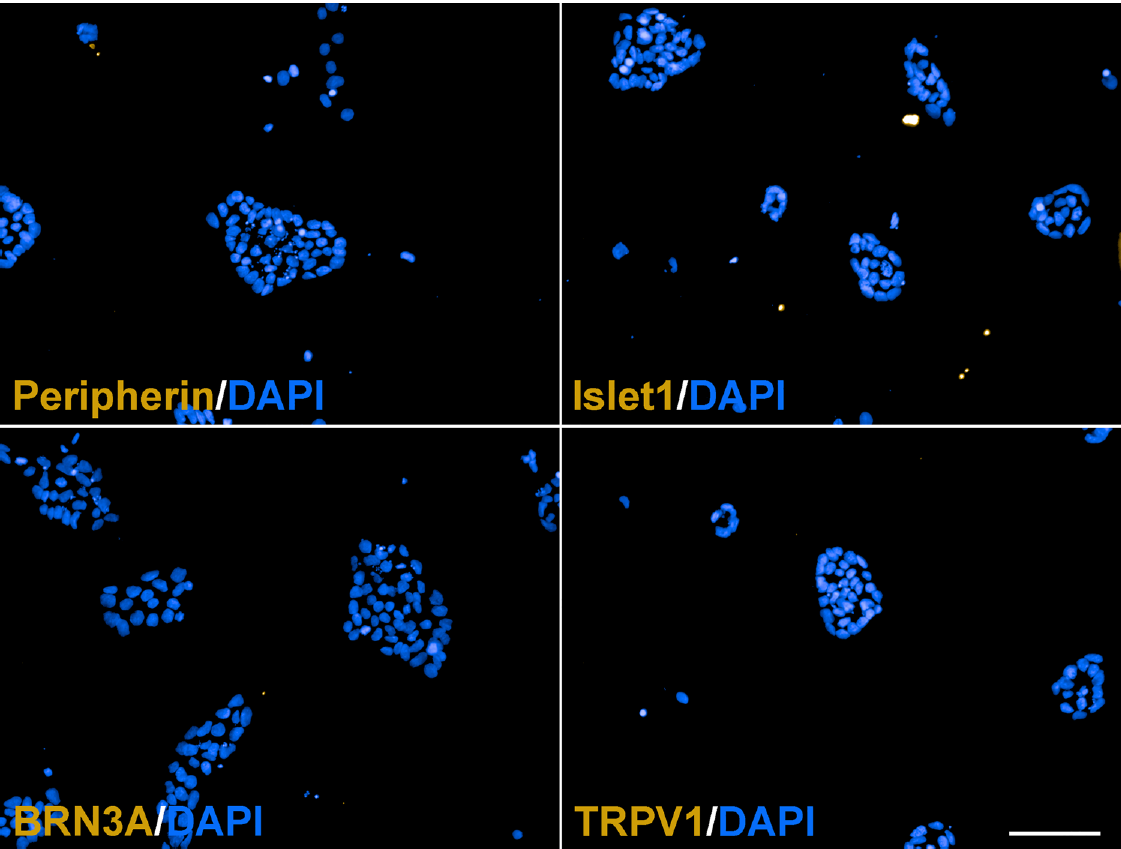
Immunofluorescence staining confirming the lack of expression of neuronal differentiation markers in the hiPSC. (A) Peripherin, (B) Islet1, (C) BRN3A and (D) TRPV1 in GM23279A hiPSC cell line. Calibration bar = 100 μm, n=2-3 independent experiments.

**Supplementary Figure 2:**
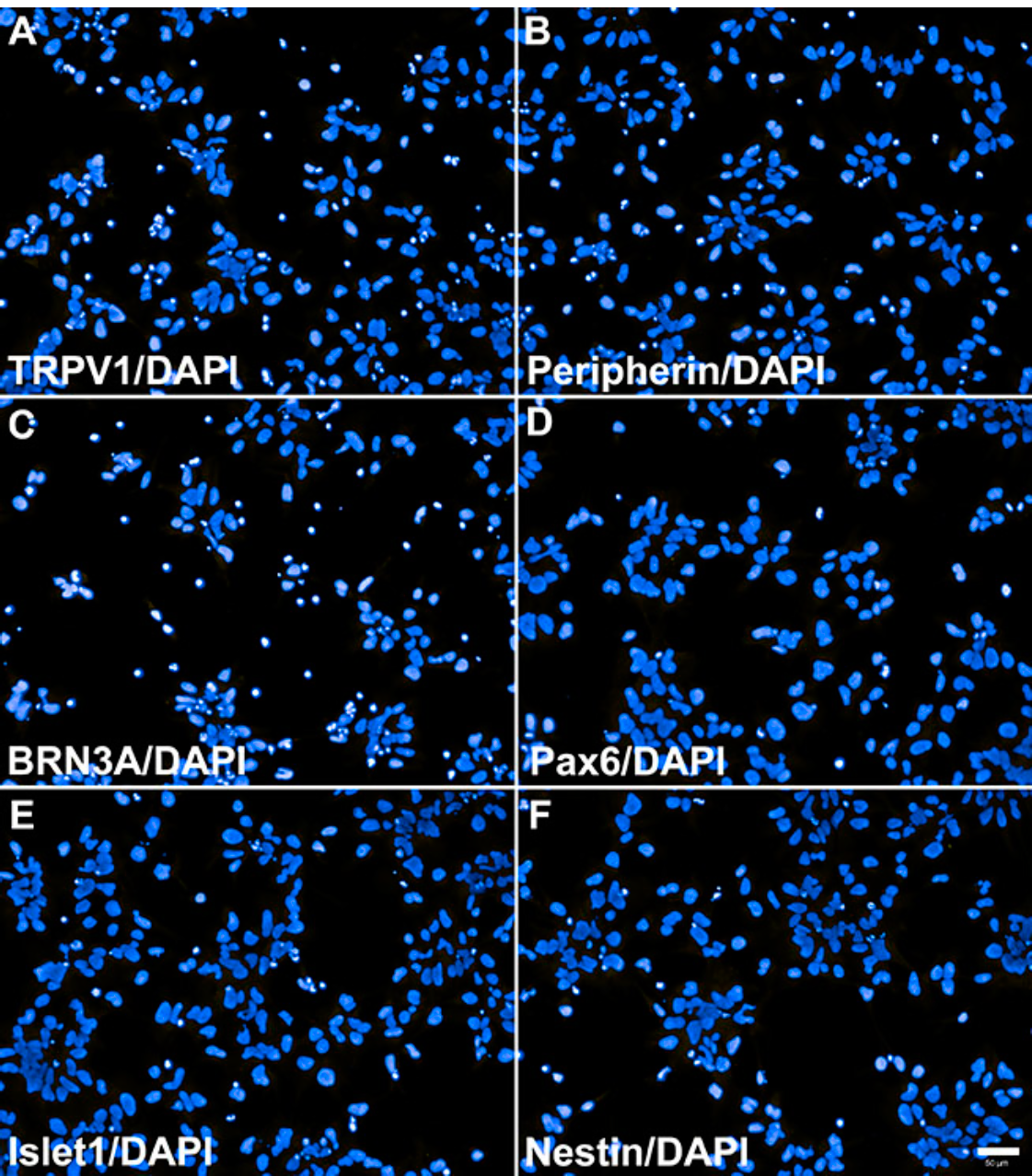
Negative controls for immunocytochemistry of NCPCs with specific markers, obtained by withholding of the primary antibody. Cells were counter-stained with DAPI for nuclei visualization. (A) TRPV1. (B) Peripherin. (C) BRN3A. (D) Pax6. (E) Islet1. (F) Nestin. Calibration bar = 100 μm, n=2-3 independent experiments.

**Supplementary Figure S3:**
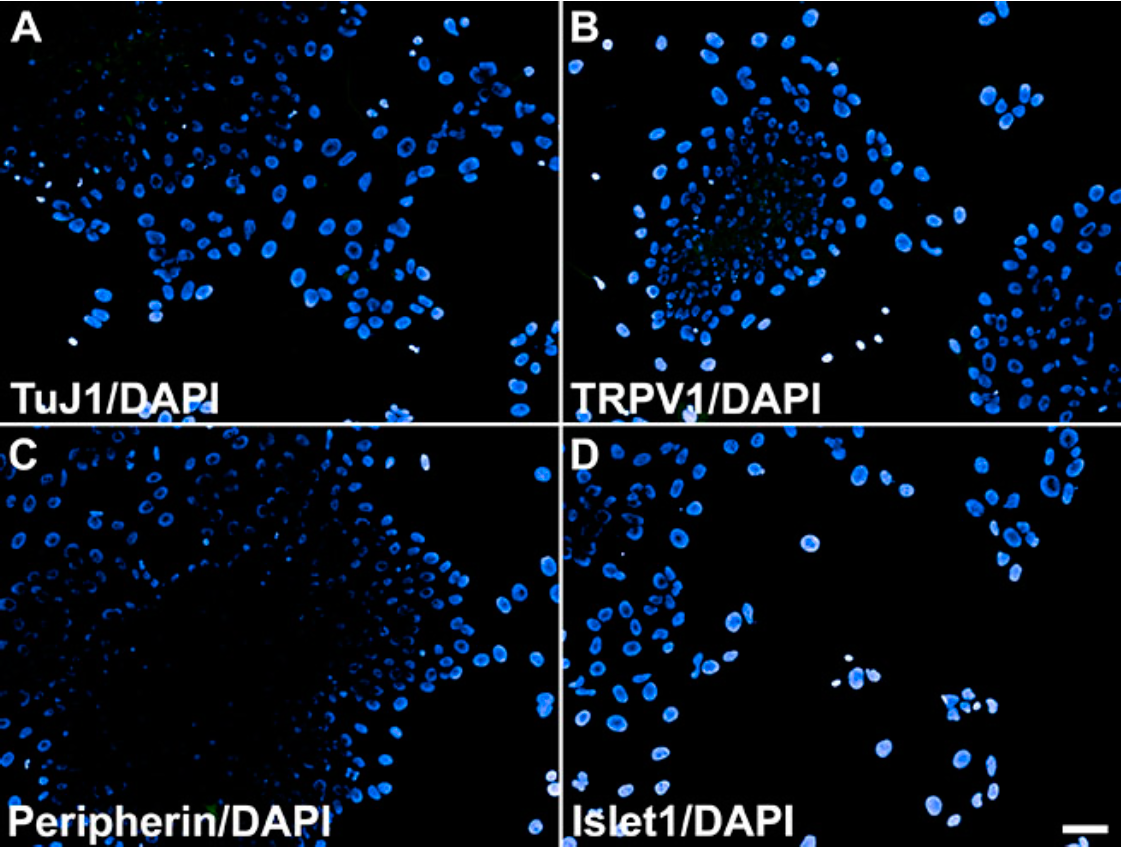
Negative controls for immunocytochemistry of peripheral sensory neurons with specific markers, obtained by withholding of the primary antibody. Cells were counter-stained with DAPI for nuclei visualization. (A) TuJ1, (B) TRPV1, (C) Peripherin, (D) Islet1. Calibration bar = 100 μm, n=2-3 independent experiments.

**Supplementary Figure S4:**
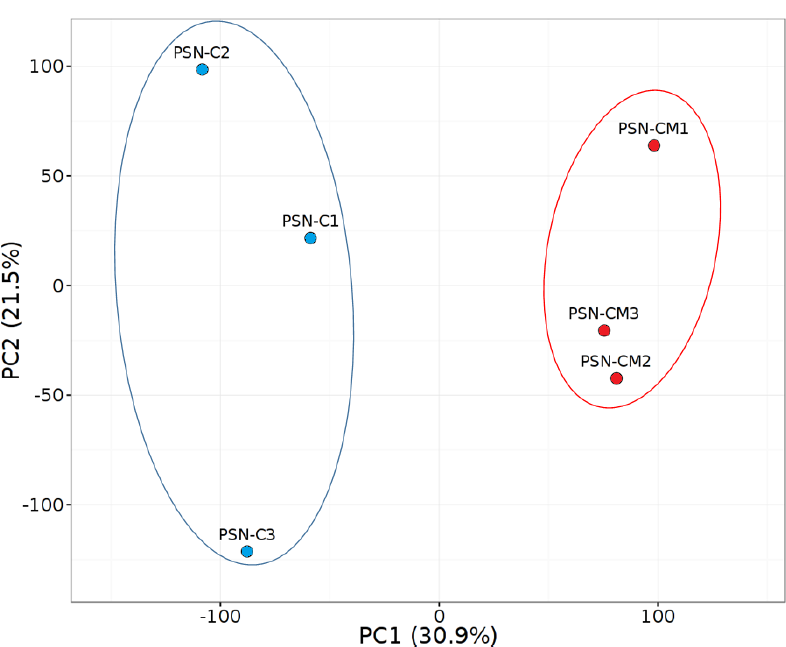
Principal component analysis of the six samples. A clear separation in expression profile between the two conditions: EP (Epilife medium) and CM (conditioned medium)showing a clear separation between the two conditions. Treatment with HEK-CM is a primary contributor to variance, and independent differentiation of cultures was a secondary component of variance. Variance between replicates was higher among HEK-CM treated cultures (CM) compared to PSN (EP) cultures.

**Supplementary Figure S5:**
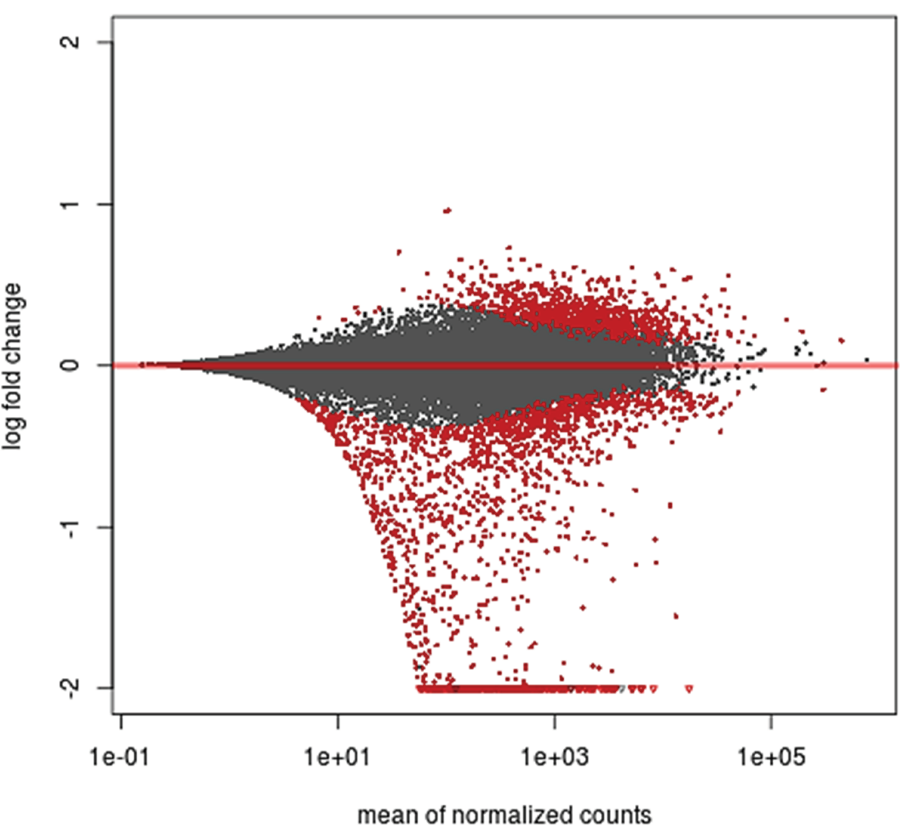
MA-plot of PSN gene expression changes induced by HEK-CM treatment. The log2 fold change for HEK-CM treatment was plotted on the y-axis and the average of the counts normalized by size factor is shown on the axis. Each gene is represented with a dot. Genes with an adjusted p value below the 0.1 threshold are shown in red. Most of the genes expressed by PSN were downregulated after the treatment with HEK-CM

**Supplementary Figure S6:**
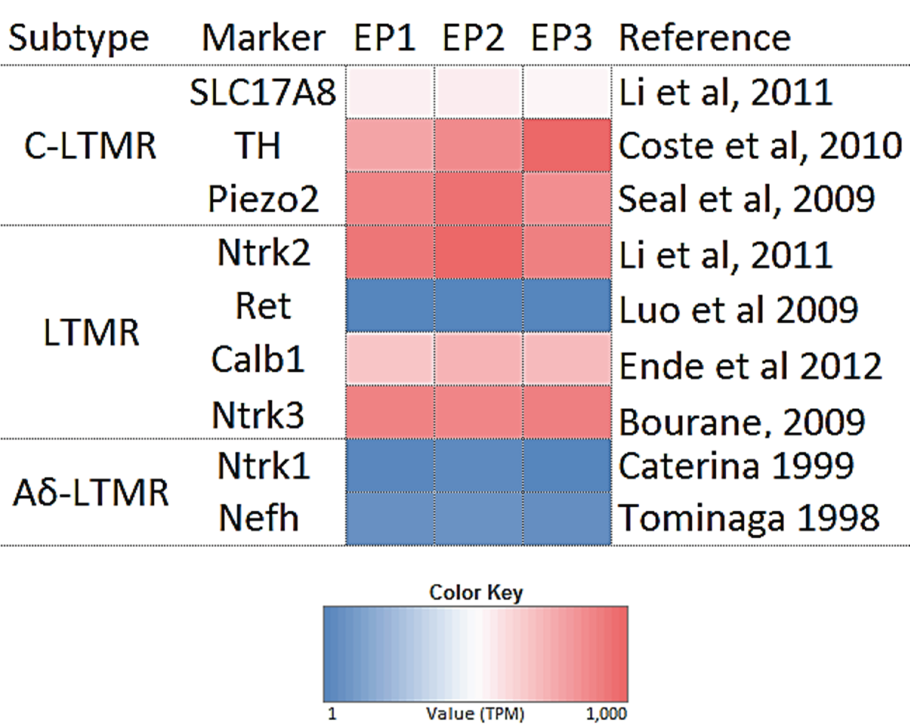
Neuronal markers expressed by hiPSC-derived PSN compared to neuronal markers reported in the literature.

**Supplementary Table 1:**
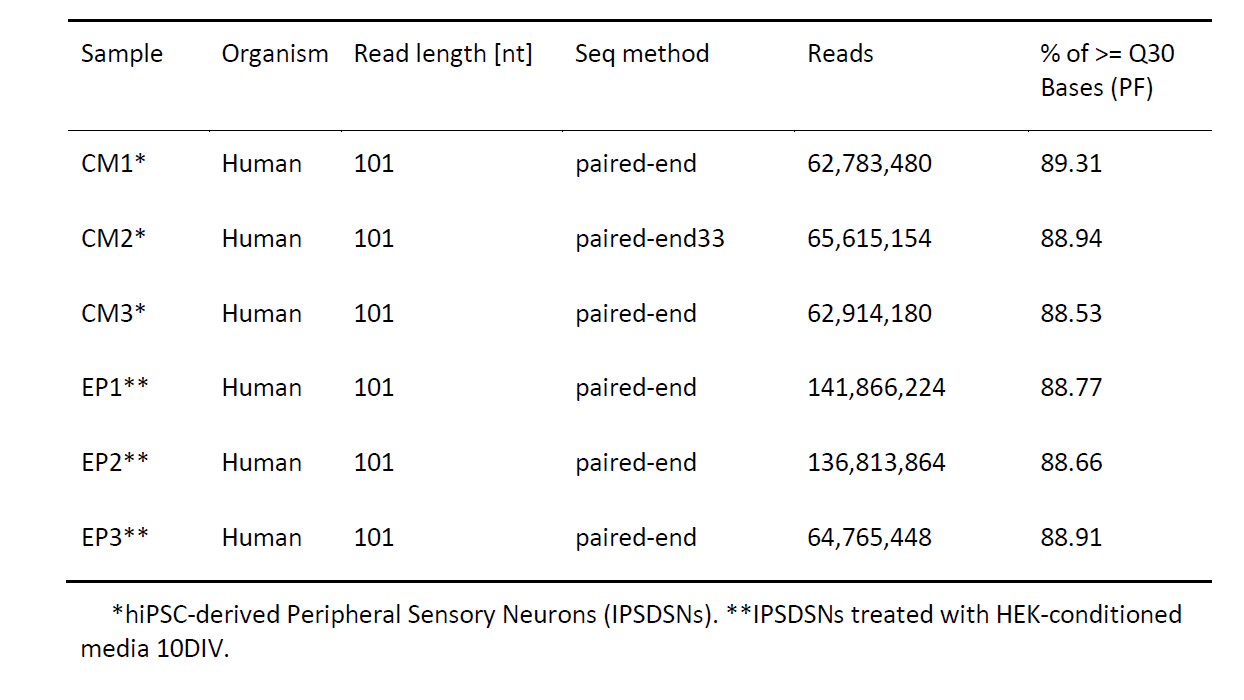
Sequencing details of RNA-Seq data-sets. Library preparation was performed for each sample to make four cDNA libraries. Each library was then sequenced with HiSeq using paired-end reads.

